# Organization of pRF size along the AP axis of the hippocampus is related to specialization for scenes

**DOI:** 10.1101/2023.05.18.541311

**Authors:** Charlotte Leferink, Jordan DeKraker, Iva Brunec, Stefan Köhler, Morris Moscovitch, Dirk B Walther

## Abstract

The hippocampus is largely recognized for its integral contributions to memory processing. By contrast, its role in perceptual processing remains less clear. Hippocampal properties differ along the anterior-posterior (AP) axis. Based on past research suggesting a gradient in the scale of features processed along the anterior-posterior extent of the hippocampus, the representations have been proposed to differ as a function of granularity along this axis. One way to quantify such granularity is with population receptive field (pRF) size measured during visual processing, which has so far received little attention. In this study, we compare the pRF sizes within the hippocampus to its activation for images of scenes versus faces. We also measure these functional properties in surrounding medial temporal lobe (MTL) structures. Consistent with past research, we find pRFs to be larger in anterior than in the posterior hippocampus. Critically, our analysis of surrounding MTL regions, such as the perirhinal cortex, entorhinal cortex and parahippocampal cortex, shows a similar relationship between scene sensitivity and larger pRF size. These findings provide conclusive evidence for a tight relationship between pRF size and the sensitivity to image content in the hippocampus.

## Introduction

It has recently become clear that visual processing triggers hippocampal activation, wherein visual representations are sourced from either external visual input or from visual information constructed from memory. As a terminus for feature-based processing along the visual hierarchy, the structure of visual receptive fields within the hippocampus and nearby medial temporal cortical regions is of key importance to understanding their role in visual perception and memory. In this study, we investigated the organization of population receptive fields (pRFs) in the hippocampus and adjacent MTL regions using a large set of high resolution retinotopy data acquired at a field strength of 7T (Benson et al., 2018). We focused on the organization of receptive field size along the anterior-posterior axis (AP) of the hippocampus and the surrounding regions that are known to project to the hippocampus: the perirhinal cortex, entorhinal cortex, and the parahippocampal cortex. As the hippocampus is stimulated by visual input and has a strong link specifically to navigation (Hirshhorn et al., 2012), we also explored the organization of pRF parameters with activation for images of scenes versus faces. As there is evidence that these stimuli are processed differently along the hippocampal long axis, as well as in the perirhinal cortex and in the surrounding MTL structures (Robin et al. 2019), we investigated the relationship of pRF size and the activation for images of scenes versus faces in these areas.

Retinotopy is a well-established characteristic of visual processing, including a contralateral representation in early visual cortical areas (Sereno et al., 1995). Recent work has shown a similar contralateral organization within the hippocampus, specifically in the middle hippocampus (Silson et al., 2021). This result gives more credence to the classification of the hippocampus as a visual processing area. Among the retinotopic mapping parameters of polar angle, eccentricity of the pRF, and size of receptive field, it has been suggested that receptive field size is related to the higher-order processing capacities in the hippocampus, such that larger pRFs are expected to relate to global and coarse-grained representations and smaller pRFs relate to fine-grained details. However, these predictions have not yet been tested using retinotopic mapping in humans yet (Evensmoen et al., 2013; Robin et al., 2016). Interestingly, this purported organization of the human hippocampus maps onto observed gradations in receptive field sizes of place cells (place fields) along the dorsoventral axis of the rodent hippocampus. These place cells generally display larger place fields in the ventral hippocampus (homologous to the human anterior hippocampus) than in the dorsal hippocampus (homologous to the human posterior hippocampus). Place cells are organized along the AP axis, analogous to humans, where fine-grained details of higher-order stimuli are processed within posterior hippocampus (Keinath et al., 2014; Kjelstrup et al., 2008; Nyberg et al., 2022), and global representations are found in the anterior hippocampus (Genon et al., 2021; Poppenk et al., 2013; Strange et al., 2014).

Representational differences within the hippocampus are suggested by differences in connectivity between anterior and posterior sections of the hippocampus with MTL cortex and other cortical regions. Coarse granularity within the anterior hippocampus is supported by the functional connectivity of the subfields located in the anterior hippocampus to the parahippocampal cortex (Dalton et al., 2019). The anterior hippocampus is responsible for processing abstract representations and is involved in tasks that require scene construction (Dalton et al., 2019; Dalton et al., 2022; Libby et al.,2012; Moscovitch et al., 2016). The posterior hippocampus is primarily connected to areas involved in visual processing, such as the retrosplenial cortex and parahippocampal cortex (Barnett et al., 2021; Dalton et al., 2022; Zeidman and Maguire, 2016). These areas are all related to scene processing broadly, which can be broken into functional objectives, such as: for the purpose of navigation or for perception (Epstein et al., 1998; Hassabis et al., 2009).

The level of spatial detail related to feature specialization found in the hippocampus activations extends to higher order representations that support functions such as spatial navigation. Learning the intricate and highly detailed information for navigating London’s city streets by car showed an increase in gray matter in the posterior hippocampus (Woollett and Maguire, 2011). Global information, such as approximating a navigation route over a long distance is processed in anterior regions during navigation and scene construction on a larger spatial scale (Brunec et al., 2018; Dalton and Maguire, 2017; Hirshhorn et al., 2012). The temporal scale for processing events also differs between the anterior and posterior hippocampus. Analogous to spatial processing capacities, the anterior hippocampus exhibits longer timescale integration whereas posterior areas show more fine-grained integration at a shorter timescale (Bouffard et al., 2021).

The current study addresses the relationship between high-level image category preferences and size of pRFs between the anterior and posterior hippocampus and the adjacent medial temporal cortex. Specifically, we ask whether the organization of size of the pRFs along the AP axis of the hippocampus and in adjacent cortical regions is congruent with the specialization for broad scene categories, where medial temporal regions have been shown to exhibit visual specialization for stimuli such as scenes, objects and faces (Barense et al., 2005; Bussey and Saksida, 2007; Murray et al., 2007; O’Neil et al., 2009, O’Neil et al., 2013).

## Methods

### Participants

We used data of 181 participants (109 females, 72 males; age 22-35) from the Human Connectome Project dataset (Van Essen et al., 2013) that were scanned at both 7 Tesla and 3 Tesla. Of the 181, we included 157 participants in our subsequent analyses (mean age = 29.5 years, 63 males, 94 females). We excluded data of 24 participants from the analysis either due to alignment failure or because we were unable to ascertain a robust scenes versus faces contrast.

### Hippocampal and MTL Segmentation

The T1-weighted anatomical images from the 3 Tesla session were processed with the HippUnfold algorithm (DeKraker et al., 2022). As a result, a Laplace gradient was calculated between the anterior (hippocampal-amygdalar transition area) and posterior (indusium griseum) endpoints of the hippocampus as described in detail in DeKraker et al. (2018). This gradient is geodesically constrained to only hippocampal grey matter and ranges from 0 (most anterior) to 1 (most posterior). This gradient is used, separately in each participant, to define coordinates in a 2D ‘unfolded’ index of points along the hippocampus. Subfields are defined according to an unfolded atlas from 3D BigBrain (Amunts et al., 2013; DeKraker et al., 2020). Laplace coordinates that determined the AP axis were binned into 3 discrete equal segments running along the geodesic anterior-posterior extent of each hippocampus in native space (Figure 1A). All gradient analyses along the continuous geodesic AP axis within the hippocampus used native space Laplace coordinates to maintain the subjects-specific curved shape of the hippocampus.

**Figure 1.**
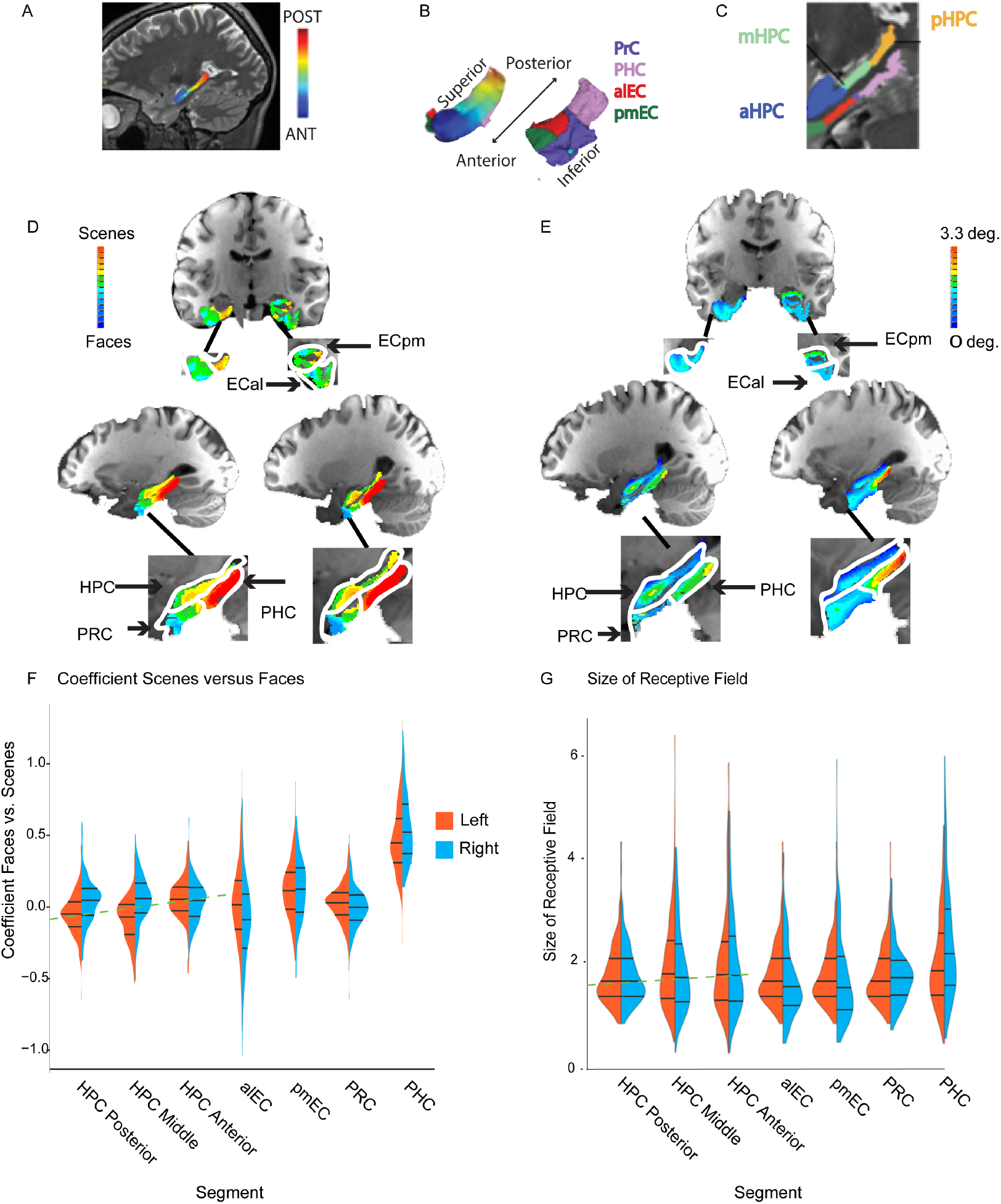
A. The gradient from anterior to posterior of the unfurled hippocampus is depicted in the saggital image on the left. B. Geodesic parcellation of the hippocampus as well as relative locations of the Entorhinal, Perirhinal, and Parahippocampal Cortex are shown in the middle. C. The right image is a close-up of the hippocampus, showing its division into three equal-sized parts. D: Average contrast coefficient for scenes versus faces depicted within the anatomical plots of the ROIs. E: Average size of receptive field depicted within anatomical plots of the ROIs. F: Quantile plots for distribution of scene vs. face coefficient for each of the 7 ROIs. G: Quantile plots for distribution of pRF size across the 7 ROIs. The positive relationship between scene selectivity and size of receptive field is evident within anterior hippocampus and parahippocampal cortex. The pattern of smaller receptive field size and less scene specialization is evident in perirhinal cortex and posterior hippocampus.

Medial temporal neocortical structures including perirhinal, entorhinal, and parahippocampal cortex were delineated using automated segmentation (ASHS) (Wisse et al., 2016). We manually segmented each subjects’ entorhinal cortex into anterolateral entorhinal cortex and postero-medial entorhinal cortex using the functional connectivity-derived protocol developed by Yeung et al., (2017), which is based on Maass et al.’s connectivity findings (2015).

### Category Specific Responses in MTL Regions

The 3T session that is part of the Human Connectome Project includes a working memory task, which is comprised of images of places, faces, tools, and body parts, taken from an assortment of studies, which focused on differences in size of stimuliper category and range of category types (Bracci et al., 2010; Kanwisher 2001; O’Craven and Kanwisher, 2000; Park and Chun, 2009; Downing et al, 2006; Peelen and Downing, 2007; Pinsk et al., 2009). Whole-brain data were collected with a 2 mm isotropic resolution and a time to repetition (TR) of 0.7s. The 3T scan session also contained a T1-weighted anatomical volume with 0.7 mm isotropic resolution. We aligned the 3T functional data to the 7T data of the same subject by computing the alignment of the 3T T1 scan to the motion-corrected 3T functional data and to the T1w scan from the 7T session. We compiled regressors for faces, places, and objects from the task stimuli files available from the HCP dataset resources. We regressed the activation for the contrast of faces versus scene to use as the coefficients to test the activation for areas that are more responsive to faces than to scenes and vice versa (3dDeconvolve in Afni). The maps were subsequently aligned to the 7T scans using an affine transform (3dAllineate in Afni) with the alignment parameters computed from aligning the anatomical scans. The quality of alignment was visually checked.

To assess the quality of the viability of the face, place, and object regressors we performed localization of the parahippocampal place area (PPA) and the occipital place area (OPA) using a (places > faces, objects) linear contrast and the fusiform face area (FFA) using a (faces > places, objects) contrast. ROIs were localized as contiguous clusters of voxels with statistically significant (p > 10-4) contrasts. These functionally localized ROIs are not used in this study, but failure to localize the ROIs (24 of 181 participants) was used as a criterion for excluding participants from the analysis.

### Population Receptive Fields

We used the retinotopy experiment performed at 7 Tesla to measure pRFs (Benson et al., 2018). The experiment consisted of 6 retinotopic runs, during which participants were shown moving bars, rotating wedges and expanding and contracting rings exposing a mélange of object, scene, and face images that are known to stimulate high-level visual cortex (Benson et al., 2018). Whole-brain functional data were collected at a 1.6 mm isotropic resolution with a repetition time (TR) of 1s. The 7T data set also included a T1-weighted anatomical scan with an isotropic resolution of 0.7 mm, which we used for data alignment with the 3-Tesla scanning session.

We converted the retinotopy data from 3dordinate space into individual subjects’ volumetric voxel space using the Neuropythy toolbox (Benson et al., 2018). We then masked the pRF data with the segmented ROI voxels (see below for ROI localization) within each participant. The pRF of each voxel was computed using the non-linear regression-based Compressive Spatial Summation algorithm (Kay et al., 2008).

### Anatomical Organization of pRFs

Differences in average size of receptive field and scene versus face contrasts across all ROIs were analyzed with a repeated-measures 2 by 7 ANOVA, of hemisphere (2 levels) by segment (7 levels).

We assessed the organization of receptive field size and scene versus face contrast coefficients along the AP axis gradient within the hippocampus using a linear mixed-effects model with participants as the random effect and location within coordinate space along the unfolded AP axis as a fixed effect to predict either size of the population receptive fields within the hippocampus or the contrast coefficient for scenes versus faces.

## Results

### Category Selectivity

Our aim was to first compare visual specialization by analyzing the coefficients from the regression analysis of neural activation for images of scenes contrasted with activation elicited by face images. To investigate the category preferences for scene versus face images within the hippocampus and in adjacent medial temporal cortex, we used a linear contrast between these types of stimuli to account for differences in absolute signal strength between and within regions of interest.

The contrast showed greater activation for scenes than faces in the hippocampus as a whole within the right hemisphere but not the left hemisphere (left hemisphere *m* = -0.026, right hemisphere *m* = 0.047). We analyzed the contrast coefficient for scenes vs. faces with a 7 × 2 rmANOVA for each of the seven ROIs and found a significant main effect for ROI segments, *F*(4.13,644.71) = 299.1, *p* < .001, and not for hemisphere, *F*(1,156) = 2.62, *p = 0.108* (Figure 1, D&F). There was a significant interaction between segment and hemisphere *F*(3.03,472.6) = 12.37, *p* < .001 with the hippocampus and surrounding regions: anterolateral entorhinal cortex; and postero-medial entorhinal cortex; parahippocampal cortex; and perirhinal cortex. Pairwise tests with Bonferroni correction to correct for multiple comparisons showed that the contrast coefficient values were significantly greater in the postero-medial entorhinal cortex than the posterior hippocampus (*p*<.001, *t* = 3.65 and 3.81 in the left and right hemispheres, respectively), and the contrast is greater for the anterior hippocampus than the anterolateral entorhinal cortex in the right hemisphere (*p* < .001, *t* = 5.95), but we found no significant difference in the left hemisphere. The perirhinal cortex showed less scene activation than the postero-medial entorhinal cortex (*p* < .001, *t* = -5.95 and -6.90 in the left and right hemispheres, respectively) and less than the middle hippocampus (*p* < .*001, t = -4.66 and* -4.31, in the left and right hemispheres, respectively). Not surprisingly, the parahippocampal cortex showed greater scene activation than all the other ROIs (*p* < .001) (Figure 1, D&F). The selectivity of the parahippocampal cortex for scenes is well documented (Epstein and Kanwisher, 1998), and is significantly stronger than for the anterolateral entorhinal cortex, postero-medial entorhinal cortex, perirhinal cortex, and even the hippocampus.

Since the magnitude of the contrast appeared to vary systematically over the length of the hippocampus, we measured the gradient of the scene versus face contrast coefficient activation along the AP axis of the unfurled hippocampus. We found that the scene advantage is greater in anterior than posterior hippocampus (Figure 1D). In fact, a mixed-effects linear regression of the face-scene contrast within each hippocampus voxel as a function of the geodesically constrained AP gradient (fixed effect) and participant (random effect) showed a significant negative slope within the left hemisphere (beta coefficient = -0.0353, standard error = 0.00101, and *t*(1191231) = 34.856,, *p* < .001,*CI* = [0.0333, 0.0373].) as well as the right hemisphere (beta weight = -0.0216, standard error 0.00102, *t*(1192400) = 21.158, *p* < .001, *CI* = [0.0196, 0.0236]).

The contrast coefficient of faces versus scenes is significant within each hemisphere, that is, the specialization for scene imagery in the hippocampus is organized in the form of a gradient along the AP axis, with greater scene specialization in anterior areas (Figure 2).

**Figure 2.**
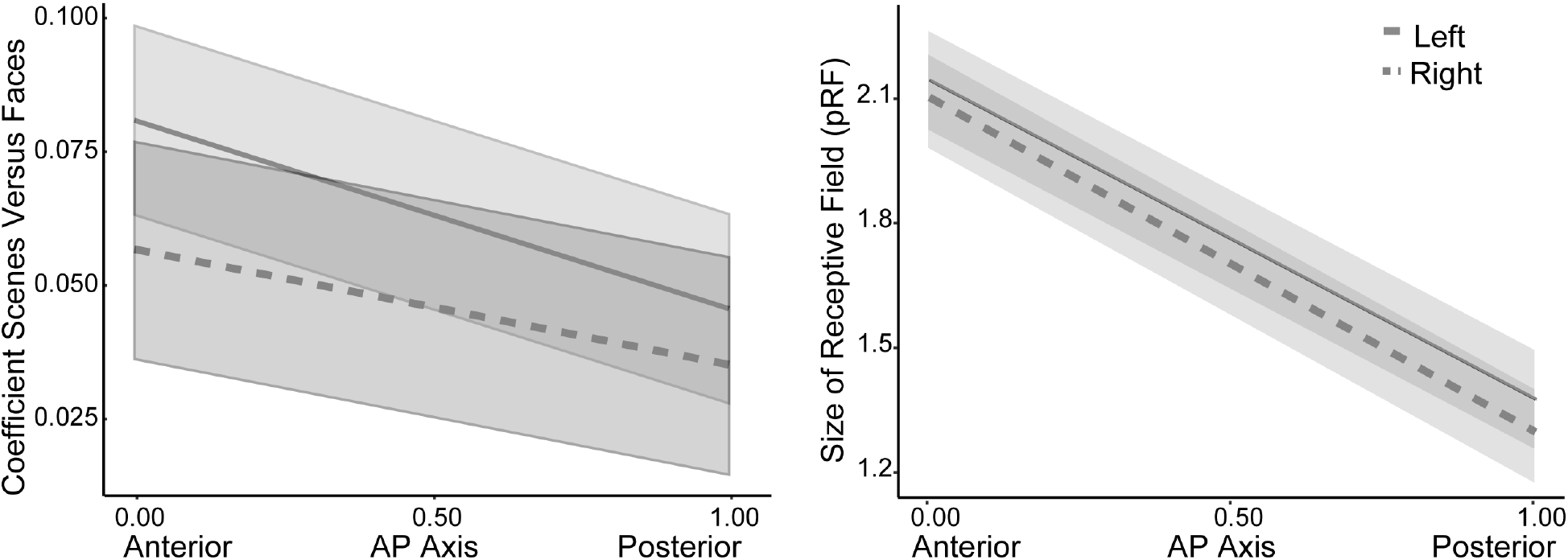
Left: The scenes versus faces contrast clearly decreases from anterior to posterior hippocampus in both the left and the right hemisphere. This slope is significant (p < .001). Right: The pRF size also varies linearly from anterior to posterior hippocampus in both hemispheres (*p* < .001), with larger receptive field size in anterior hippocampus.

### Size of Population Receptive Fields

We compared the size of pRFs in medial temporal cortex regions surrounding hippocampus that are known to have direct inputs to the hippocampus: the entorhinal cortex, the parahippocampal cortex, and the perirhinal cortex. Our analysis of pRF size (sigma of the fitted 2D Gaussian) via a 7 × 2 rmANOVA yielded a significant main effect for segments, *F*(4.28, 667.82) = 26.29, *p* < .001, but not for hemisphere, *F*(1, 156)= 0.48, *p = 0.49* (after Greenhouse-Geisser correction for sphericity). There was a significant interaction between segment and hemisphere *F*(4.96, *774.22*) = 6.32 *p* < .001 in the hippocampus and surrounding ROIs. Post-hoc analyses, which were conducted using the Bonferroni method to correct for multiple comparisons, showed that the activation levels were different between parahippocampal cortex and all other segments, *p* <.001 (Figure 1, E&G). Within the hippocampus, voxels in the anterior and middle segments had significantly larger receptive fields than those found in the posterior hippocampus regions (*p* < .001). The middle hippocampus and anterior hippocampus had larger receptive fields than the perirhinal cortex, combined across hemispheres (*p* = .0018 and *p* < .001, respectively). The middle and anterior hippocampus also had larger pRFs than the postero-medial entorhinal cortex and anterolateral entorhinal cortex averages, combined across hemispheres.

We measured the average size of population receptive fields for each voxel within the hippocampus as well as adjacent cortical regions (Figure 1E). Within the hippocampus, we find a clear gradient along the AP axis with significantly larger pRFs in anterior regions than in the posterior hippocampal regions (Figure 2): within the left hemisphere, the mixed-effects regression with AP axis location as a fixed effect and participants as a random effect showed a significant negative relationship *t*(1190000) = -83.13, *beta* = -0.7678, *p* < .001, *sem* = 0.00924, *CI* = [-0.7859, -0.7497]. The right hemisphere showed a similar significant negative relationship, *t*(1192000)=-85.58, *p* < .001, *beta* = -0.8054, *sem* = 0.009411, *CI* = [-0.8238, -0.787].

### Relationship Between Category Selectivity and Population Receptive Field Size

The larger population receptive fields in the anterior compared to the posterior hippocampus align well with the stronger preferences for scenes in the anterior compared to the posterior hippocampus. We tested this relationship formally with a mixed-effects regression model with contrast coefficient as the predicted variable, pRF size of hippocampus voxels as a fixed effect and participants as a random effect. We found a significant relationship with pRF size in the left hippocampus, *p* < .001, *t*(1191000) = 25.224, *stderror* = 0.0001, *beta* = 0.02525, as well as the right hippocampus, *p* < .001, *t*(1192000) = 11.005, *stderror* = 0.0001, *beta* = 0.001. That is, scenes versus faces selectivity is strongly related to population receptive field size in both hemispheres.

The contrast coefficient differences for scenes vs. faces across the medial temporal ROIs show a similar increase of the average size of receptive fields across hippocampus segments, where generally smaller pRFs were related to face processing, and larger pRFs showed greater activation for scenes. To test this relationship explicitly, we performed the same mixed-effects regression analysis as above, separately for the voxels within each of the cortical ROIs. Our results of a mixed effects regression with participants as the random factor and size of receptive field as the independent variable, showed that the scene vs. face contrast coefficients are significantly related to size of receptive field, bilaterally, within parahippocampal cortex (left: *p* < .001, *t*(435500) = 52.42, *stderror*=0.0098, *beta* = 0.515; right: *p* < .001, *t*(432500) = 81.45, stderror = 0.00925, *beta* = 0.753). Within the right perirhinal cortex, we found a significant relationship of larger receptive field size with greater scene activation (*p* < .001, *t*(1172000) = 24.86, *stderror* = 0.0051, *beta* = 0.127), and within the left perirhinal cortex, a negative relationship between pRF size and scene vs. face contrast coefficient (*p* < 0.001, *t*(1201000) = -15.05, *stderror* = 0.0052, *beta* = -0.0786). There was no consistent pattern of relationship with pRF size and scene selectivity within postero-medial entorhinal cortex (left hemisphere was non-significant, *p* = 0.213, *t*(111500)=-1.246, *stderror* = 0.0161, *beta* = -0.021; right hemisphere: *p* < .001, *t*(95040)=9.346, *stderror* = 0.0173, *beta* = 0.162). The relationship within anterolateral entorhinal cortex was also inconsistent across hemispheres (left: *p* < .001, *t*(121200) = 12.50, *stderror* = 0.0132, *beta* = 0.165; right: *p* < .001, *t(*128100) = -7.065, *stderror* = 0.0123, *beta* = -0.0869).

## Discussion

Our findings generally confirmed our prediction that pRF size would vary along the longitudinal axis of the hippocampus, decreasing from anterior to posterior. We also found pRF size to be related to specialization of processing higher order stimuli, with faces preferentially processed in the posterior hippocampus with its relatively small pRFs, and scenes being preferentially processed in the anterior hippocampus with its larger pRFs (Figure 2). Larger pRF size in the anterior hippocampus complements research showing that the scale of feature-based processing varies along the AP axis, with more fine-grained, feature-based processing occurring in posterior and more schematic, and therefore larger scale, processing occurring in the anterior hippocampus (Poppenk et al., 2013; Brunec et al., 2018).

By and large, we found similar relations between the size of pRFs and specialization of stimulus processing in the MTL regions surrounding the hippocampus. Namely, the scene-selective parahippocampal cortex showed significantly larger pRF sizes than the less scene-selective areas, namely: the perirhinal cortex, and the entorhinal cortex (Arcaro et al., 2009).

Specialization for perceptual processing of scene or face images in high-level visual cortex is a well-documented phenomenon. The hippocampus is integral to memory creation and is not often characterized as a visual perception region. Here, we observed a similar effect size for large receptive fields and scene processing within the anterior hippocampus as we did in the parahippocampal cortex. Our results showed activation to be higher for scenes in the anterior hippocampus and higher for faces in the posterior hippocampus (Figure 1D), which supports the theory that the hippocampus is involved in perceptual processing of high-level visual features.

Our findings also support the hypothesis that larger, more global features and schematic representations activate the anterior hippocampus during visual processing (Audrain et al., 2022; Farzanfar et al., 2023), consistent with the view that the anterior hippocampus specializes in scene construction (Dalton and Maguire,2017).

Category preference in perirhinal cortex aligns with lesion studies specifically pointing towards its importance in object recognition (Barense et al., 2005). Previous fMRI research on perirhinal cortex showed greater activation for faces than for scenes, where visual representations are products of complex visual features (Bussey and Saksida, 2007; Murray et al., 2007; O’Neil et al., 2009, O’Neil et al., 2013). The face processing network also showed connectivity between perirhinal cortex and FFA (O’Neil et al., 2014).

The anterolateral entorhinal cortex showed greater face activation than the postero-medial entorhinal cortex, as well as relatively smaller pRFs. This finding is consistent with literature showing greater connectivity of perirhinal cortex to anterolateral than postero-medial entorhinal cortex, especially in the right hemisphere (Maass et al., 2015; Ferko et al., 2022). The fine-grained processing requirements for face recognition align with the smaller average pRF size in perirhinal cortex as well as in the anterolateral entorhinal cortex.

In contrast to the hippocampus, the postero-medial entorhinal cortex did not show a particularly strong preference for faces or scenes, nor was the size of receptive fields in this region related to perceptual specialization for scenes or faces. This deviation from the intuitive relationship of pRF size and feature specialization could be explained by the spatial integration that is thought to occur in postero-medial entorhinal cortex (Avidan and Behrmann, 2021). It is a region that is less involved in feature processing but involved in combining high-level feature maps within the MTL regions for organization of location information in physical space.

We expected to find larger receptive fields in parahippocampal cortex given past literature showing that large pRFs are a characteristic of perceptual scene processing areas (Arcaro et al., 2009). The large pRF size supports the theory that scene images tend to be processed more globally (Oliva and Torralba, 2006). We also expected to find larger pRFs in the anterior hippocampus, although it is typically not considered to be a perceptual processing area due to its specialization for processing spatial schemas, and therefore, associated with abstracted representations from memory (Poppenk et al., 2013; Steel et al., 2020).

Our findings revealed a closer relationship of pRFs with visual activation, suggesting that the closer relationship is linked more strongly to perceptual processing. Although contralateral hemifield retinotopy, which is a characteristic of perceptual processing areas, has been found in the hippocampus (Silson et al., 2021), those results were inconsistent across all three segments of the hippocampus. This inconsistency in finding a hemifield bias could reflect a weaker involvement of the hippocampus in the processing of visual features. Specialization within the hippocampus for visual perception of scenes versus faces was evident in a systematic review that showed inconsistent activation between the anterior and posterior parts of the hippocampus (Robin et al., 2019). Our results of greater scene image activation within anterior and middle hippocampus align with most studies that used the scenes versus face contrast, as men-Honed in Robin et al. (2019). The discrepancy across studies may be specific to the one-back task that required greater memory recall instead of the detailed spatial navigation that has been shown to recruit the posterior hippocampus (Hirshhorn et al., 2012; Woollett et. al, 2011).

Our results of larger pRFs in the anterior hippocampus seem to conflict with the retinotopic connectivity of V1 with the anterior hippocampus during movie watching (Knapen, 2021). Stronger activation of the anterior hippocampus and, as a consequence, stronger functional connectivity during movie viewing than for resting state could be explained by the movies’ story narrative that recruits the need for the representation of longer timeframes, which the anterior hippocampus supports, as shown by Brunec et al. (2018) and Bouffard et al. (2022). This interpretation would lead to important questions for future research about how temporal scale interacts with spatial scale in the hippocampus.

## Conclusions

Visual specialization in the hippocampus is known to vary along the anterior-posterior axis, where anterior areas have a global bias, and posterior areas are more involved in fine-grained processing. Here, we found that visual specialization of parts of the hippocampus tends to mirror the size of population receptive fields, where greater specialization of the anterior hippocampus for scenes was associated with larger pRFs and greater specialization of the posterior hippocampus for faces was associated with smaller pRFs.

This organization of pRF size along the AP axis along with the specialization for particular image content within areas of the hippocampus, overlap with mnemonic processing of visual information. Our findings suggest that pRF representations in the hippocampus and surrounding medial temporal regions are indicative of high-level and abstract representations that extend beyond direct perceptual presentations.

## Acknowledgments

We would like to thank to Kayla Ferko and Alexander Minos for the manual segmentation work on the entorhinal cortex and to Claudia Damiano for help with processing the 3T data for the scene-face contrast.

## Supplementary Material

**Supplementary Figure 1.**
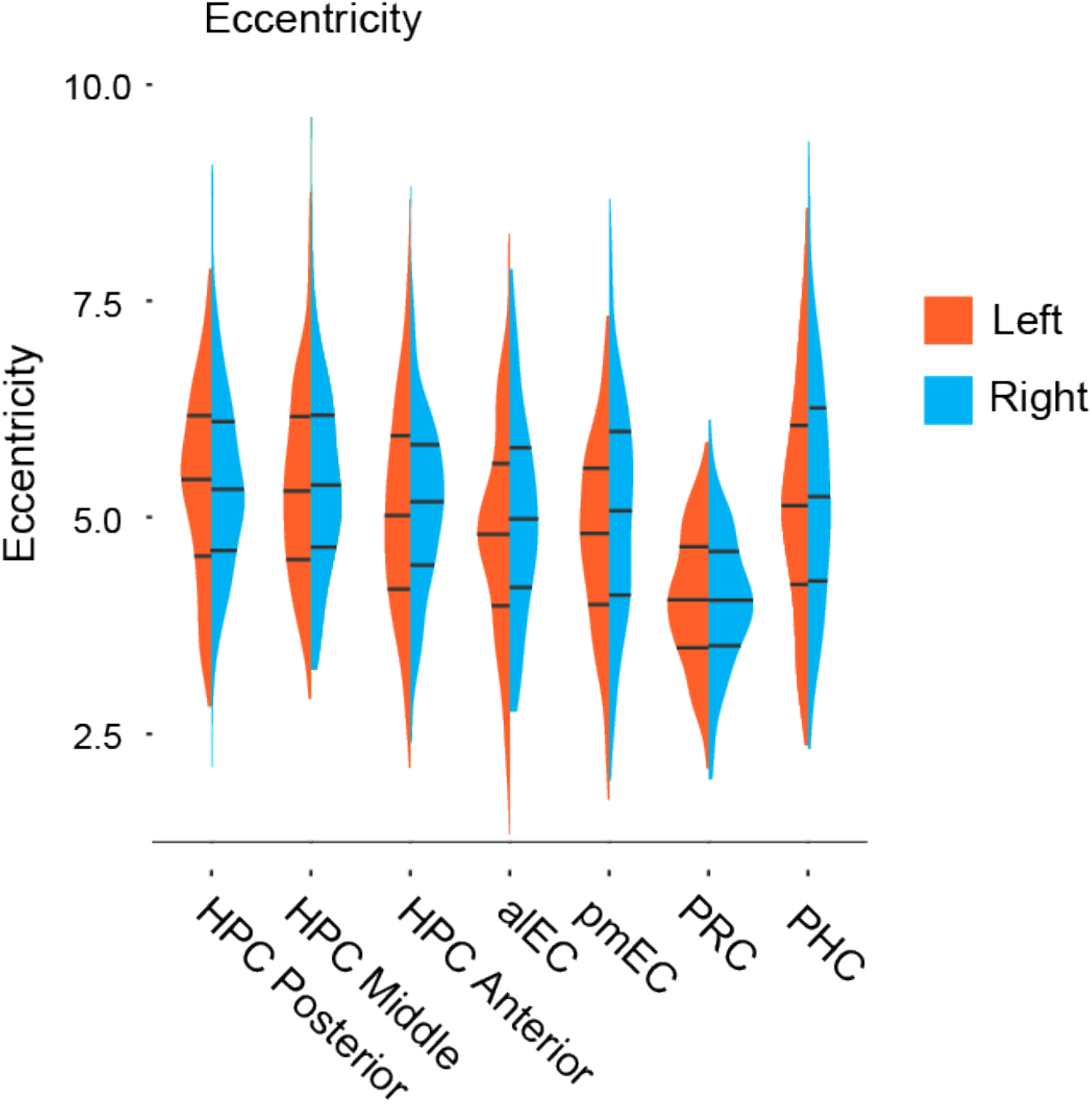
*Average Eccentricity across surrounding segments of the* hippocampus. A repeated-measures ANOVA for eccentricity parameter by segment yielded a main effect for segment, F(4.98,776.52) = 52.77, p < .001, and for hemisphere, *F*(1,156)= 6.23, *p=.014*. There was not a significant interaction between segment and hemisphere, *F*(4.96,773.48)=1.37 *p* = .23.

